# High-resolution repertoire analysis of Tfr and Tfh cells reveals unexpectedly high diversities indicating a bystander activation of follicular T cells

**DOI:** 10.1101/231977

**Authors:** Paul-Gydéon Ritvo, Wahiba Chaara, Karim El Soufi, Benjamin Bonnet, Adrien Six, Encarnita Mariotti-Ferrandiz, David Klatzmann

**Affiliations:** Sorbonne Université, INSERM, UMR S 959, Immunology-Immunopathology-Immunotherapy (I3); F-75005, Paris, France; Biotherapy (CIC-BTi) and Inflammation-Immunopathology-Biotherapy Department (DHU-i2B), Hôpital Pitié-Salpêtrière, AP-HP, F-75651, Paris, France

**Author notes:** Corresponding author: David Klatzmann.

## Abstract

T follicular helper (Tfh) and regulatory (Tfr) cells regulate B cell activation and ultimately antibody production. While concordant results show that Tfh cells are specific for the immunizing antigens, limited and even controversial results have been reported regarding the specificity of Tfr cells. Here we used high-throughput T cell receptor (TCR) sequencing to address this issue. We observed that although the Tfh- and Tfr-cell repertoires are less diverse than those of effector (Teff) and regulatory T (Treg) cells, they still represent thousands of clonotypes after immunization with a single antigen. T-cell receptor beta variable (TRBV) gene usage distinguishes both follicular T cells (Tfol) from non-Tfol cells, as well as helper (Teff and Tfh) vs. regulatory (Treg and Tfr) cells. Analysis of the sharing of clonotypes between samples revealed that a specific response to the immunizing antigen can only be detected in Tfh cells immunized with a non-self-antigen and Tfr cells immunized with a self-antigen. Finally, the Tfr TCR repertoire is more similar to that of Tregs than to that of Tfh or Teff cells. Altogether, our results highlight a bystander Tfol-cell activation during antigenic response in the germinal centres and support the Treg cell origin of Tfr cells.

**Significance Statement:** Follicular helper T (Tfh) cells promote high-affinity antibody production by B cells while follicular regulatory T (Tfr) cells represses it. The question of the specificity of follicular T (Tfol) cells is of utmost importance in the understanding of the antibody response specificity and our work is the first to analysed the global Tfol TCR repertoire in wild type mice. This allowed us not only to portray the overall global structure of these repertoires, but also to substantiate the fact that Tfr cells respond to self-antigen while Tfh cells respond to non self-antigen, a still controversial issue. Importantly, our work revealed an unexpected bystander activation of Tfol cells. We think and discuss that it has a general significance in immune responses and possibly immunopathologies.

## INTRODUCTION

The germinal centre (GC) is an essential structure for the activation of B cells and the generation of high-affinity antibodies providing a humoral protection against pathogens (1, 2). Follicular helper (Tfh) cells promote the differentiation and activation of B cells into plasma cells, enabling antibody production (3, 4). In contrast, the recently discovered T follicular regulatory (Tfr) cells (5–7) inhibit this GC reaction, therefore reducing antibody production, in humans(6) and mice (7). Tfr also promote high-affinity antibodies, as mainly low-affinity antibodies were detected in the absence of Tfr (8, 9). In addition, Tfr are increased in infectious diseases with insufficient antibody production (10–13). Contradictory results have been found so far in autoimmune diseases, where Tfr are either increased or decreased (14–17). Their suppressive effect on Tfh and GC B cells is associated with the expression of CTLA-4 (18, 19). We recently showed that Tfr control Tfh by IL-1 deprivation (20). This unknown mechanism was revealed after redefining Tfr phenotype as CD4^+^CXCR5^hi^PD1^hi^ Foxp3^−^CD25^−^ (20), as reported by others (21). Most studies on Tfr were performed on CD4^+^CXCR5^+^PD1^+^Foxp3^+^CD25^+^ (22) cells, which we found to be enriched in conventional regulatory T cells (20). Therefore, those recent observations highlight the need to revisit many aspects of Tfr cell biology as previous studies might have reported results from a mixture of Tfr cells and regulatory T cells (Treg cells) (20).

Two such aspects are Tfr specificity and origin. It is believed that to activate B cells, Tfh cells must recognize the same antigen as the B cells they help (23, 24). Several studies based on peptide-class II major histocompatibility complex (pMHC-II) tetramers (25, 26) or ELISPOT assays (27) established that Tfh cells are specific for peptides of the immunizing antigen. However, contradictory results have been obtained for Tfr cells. A recent study addressed this issue by comparing the T cell repertoires after immunization with the myelin oligodendrocyte glycoprotein (MOG) in wild-type and MOG knock-out mice. MOG is a self-antigen in the former and a non-self antigen in the latter. In both conditions, Tfr specific for the immunodominant peptide of MOG assessed by tetramers were identified, indicating that independently of the self or foreign nature of the immunizing antigen, Tfr cells could be specific for it (28). They also showed that Tfh cells share with Tfr cells TCRs specific for the immunizing antigen (28), suggesting that Tfr cells can differentiate from naïve helper T cells. In contrast, in a previous study using high-throughput sequencing (HTS) in immunized mice with a fixed TCR β chain, we reported oligoclonal expansion in Tfh cells and a broad TCR usage in Tfr cells from the same GCs (29). In addition, only Tfh specific for the immunizing antigen were found in draining lymph nodes, not Tfrs, and the Tfr repertoire was closer to Treg than to Tfh, suggesting a Treg origin. However, all these results (28, 29) were obtained with CD4^+^CXCR5^hi^PD1^hi^Foxp3^+^CD25^+^ “Tfr” cells. Therefore, the observed TCR repertoire diversity and composition was probably that of a mixture of TCRs derived from Treg and Tfr cells.

In these previous studies, it should be noted that Tfr cell specificity was studied either (i) with tetramers that assess only limited specificities (28) or (ii) using HTS in TCR-transgenic mice with a biased repertoire (29). Here, we performed HTS on stringently defined Tfr cells CD4^+^PD1^hi^CXCR5^hi^Foxp3^+^CD25^−^ from wild-type mice and compared their TCR repertoires to that of Tfh, Treg and Teff cells by HTS targeting the β chain. Cells were obtained from nonimmunized mice or mice immunized either with a self-antigen (insulin, INS) or a non-self-antigen (ovalbumin, OVA). Our work reveals an unexpected high diversity of the repertoire of Tfol cells indicative of a bystander activation during antigen-specific responses and support a Treg cell origin for Tfr cells.

## RESULTS

To investigate the TCRβ repertoire of Tfr cells, we used mice with the transgenic expression of green-fluorescent protein (GFP) under the promoter of Foxp3 to purify Tfh (CD4^+^CD8^−^ CXCR5^hi^PD1^hi^Foxp3^−^), Tfr (CD4^+^CD8^−^CXCR5^hi^PD1^hi^CD25^−^Foxp3+), Teff (CD4+CD8^−^CXCR5^lo/−^PD1^lo/−^ Foxp3^−^) and Treg (CD4^+^CD8^−^CXCR5^lo/−^PD1^lo/−^Foxp3^+^) cells. In order to recover sufficient amounts of the scarce Tfr-cell population, cells were purified from pools of six to eight mice (later named “individuals”) receiving the same treatment. We generated three pools of mice immunized intraperitoneally with OVA in alum, three pools of mice immunized with insulin (INS) in alum and two pools that were not immunized in order to study cells at homeostasis. Finally, we massively sequenced T-cell mRNA from sorted Tfr-, Tfh-, Teff- and Treg-cell subsets

### Tfol cells have a lower diversity than non-Tfol cells

We first analyzed the TCR diversity of the four cell subsets in the three immunization settings. Sequence and clonotype numbers obtained for all samples are provided in **Supplementary Table S1** for reference. Rarefaction curves (**Fig. 1A**), which represent the observed numbers of different clonotypes (i.e. unique combination of TRBV-CDR3p-TRBJ) observed at a given sample size (i.e. number of TR sequences), showed that both Tfr- and Tfh-cell repertoires are less diverse than those of Teff and Treg cells. Indeed, from 100 000 sequences and over, Tfol-cell clonotype numbers are systematically two-to three-fold lower than that of Teff/Treg cells (p=0.0079, Mann-Whitney U). At the sequencing depth used, we have almost captured the entire diversity of Tfh and Tfr cell TCR repertoires, as shown by the almost flat slopes, while this is not the case for Teff and Treg cells.

**Fig. 1:**
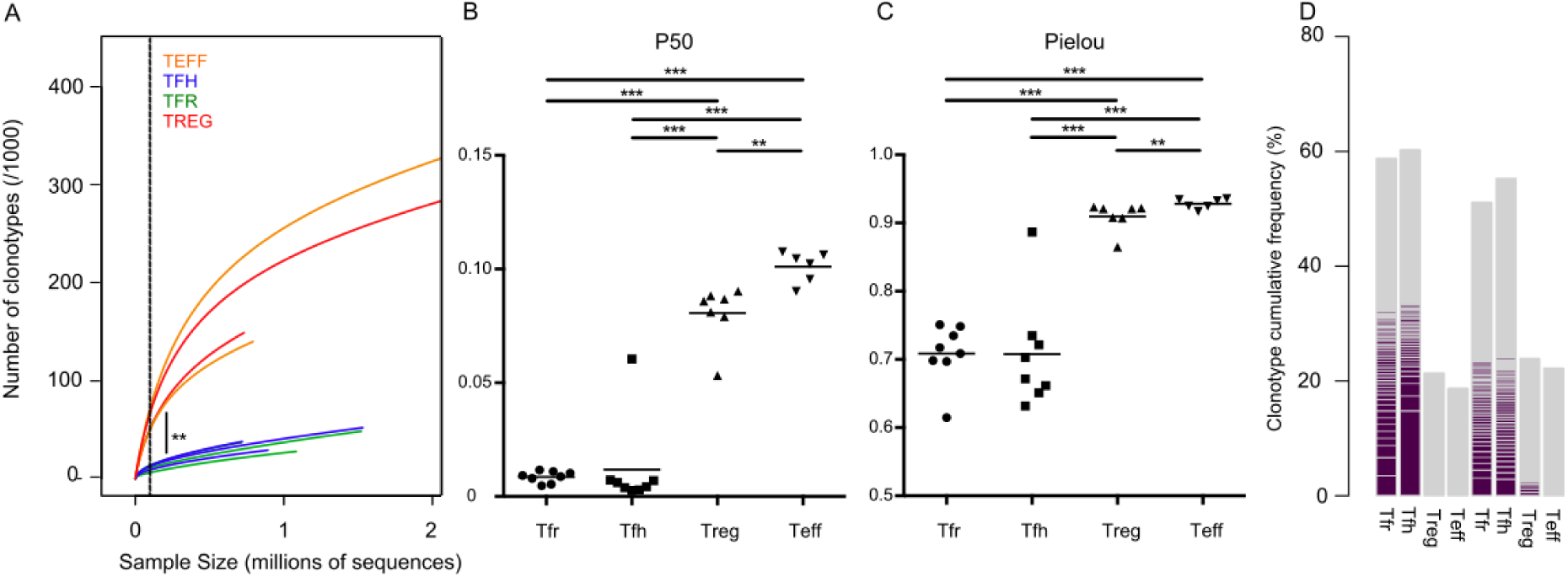
Tfol cells display a lower diversity than non-Tfol cells, yet are polyclonal. (**A**)Representative rarefaction curves displaying the number of clonotypes as a function of the number of reads (sample size) were computed for two samples of each T cell subset randomly selected from OVA-immunized mice. For a sample size of 100 000 (vertical line), the number of unique clonotypes is higher in Treg and Teff cells than in Tfr and Tfh cells (p=0.0079, MannWhitney U). (**B-C**) P50 (**B**) and Pielou’s evenness (**C**) indices were calculated for all the samples of each T cell subset (see **Supplemental Table 1** for details) and compared by the MannWhitney U test (*p<0.05; **p<0.01, ***p<0.001). (**D**) Cumulative frequencies of the 1% predominant clonotypes for each of the 4 T cell subsets were calculated. One histogram bar represents one sample and one coloured line one clonotype.

Other commonly used diversity indices confirmed these observations. The percentage of the most predominant clonotypes that account for 50% of the sequences of a sample size (P50) of Tfol cells (1 to 2%) is significantly lower than the average P50 of Teff and Treg cell repertoires (**Fig. 1B**). This marked difference between Tfol and non-Tfol cells, which is more pronounced than the one evidenced by rarefaction curves, suggests that there are important clonotype expansions in Tfol cells. The Pielou index(30), ranging from 0 to 1, assesses the evenness of a repertoire: the higher the index the more equally represented the clonotypes are. Repertoire evenness is both high and yet significantly different for Treg and Teff cells. In contrast, the Pielou index is significantly lower for Tfol cells compared to non-Tfol cells (**Fig. 1C**). The lower richness described by the rarefaction curves and the P50 index could thus be explained by a higher number of expansions among Tfr and Tfh cells compared with Teff and Treg cells.

This was confirmed by analyzing the frequency of the predominant clonotypes in the different cell populations (**Fig. 1D**). The predominant 1% of clonotypes for Tfr and Tfh cells represent at least 50% of the total repertoire, compared to only 10% for Teff and Treg cells. Some clonotypes represent more than 1% of the sample in Tfr- and Tfh-cell samples, which is not observed in Treg and Teff cells. Finally, reconstruction of immunoscope profiles from NGS data showed a Gaussian distribution profile for Teff and Treg cells, and numerous expansions in Tfol-cell samples, as exemplified for representative TRBVs (**Supplementary Fig. S1**). Altogether, these results suggest that the global characteristics of the Tfh- and Tfr-cell repertoires are similar and that both subsets have a skewed diversity compared to non-Tfol cells.

### TRBV but not TRBJ genes are differentially used by the four cell populations

We performed a principal component analysis (PCA) of TRBV (**Fig. 2A**) and TRBJ (**Fig. 2B**) gene frequencies. Strikingly, the first component (PC1) of the PCA using TRBV gene frequencies (**Fig. 2A**), which explains about ≈32% of the variability, separates very well Tfol-from non-Tfol-cell subsets, while the second (PC2; 12% of the variability) separates the regulatory subsets (Tfr and Treg cells) from the non-regulatory ones. In contrast, no clear separation of the four T cell subsets (**Fig. 2B**) could be observed using TRBJ gene usage. The same analysis for the TRBVBJ combination frequencies (**Fig. 2C**) showed clear separation of the four T cell subsets, although only ≈30% of the variability is explained by the first two components compared to ≈50% when focusing on the TRBV gene frequencies only.

**Fig. 2:**
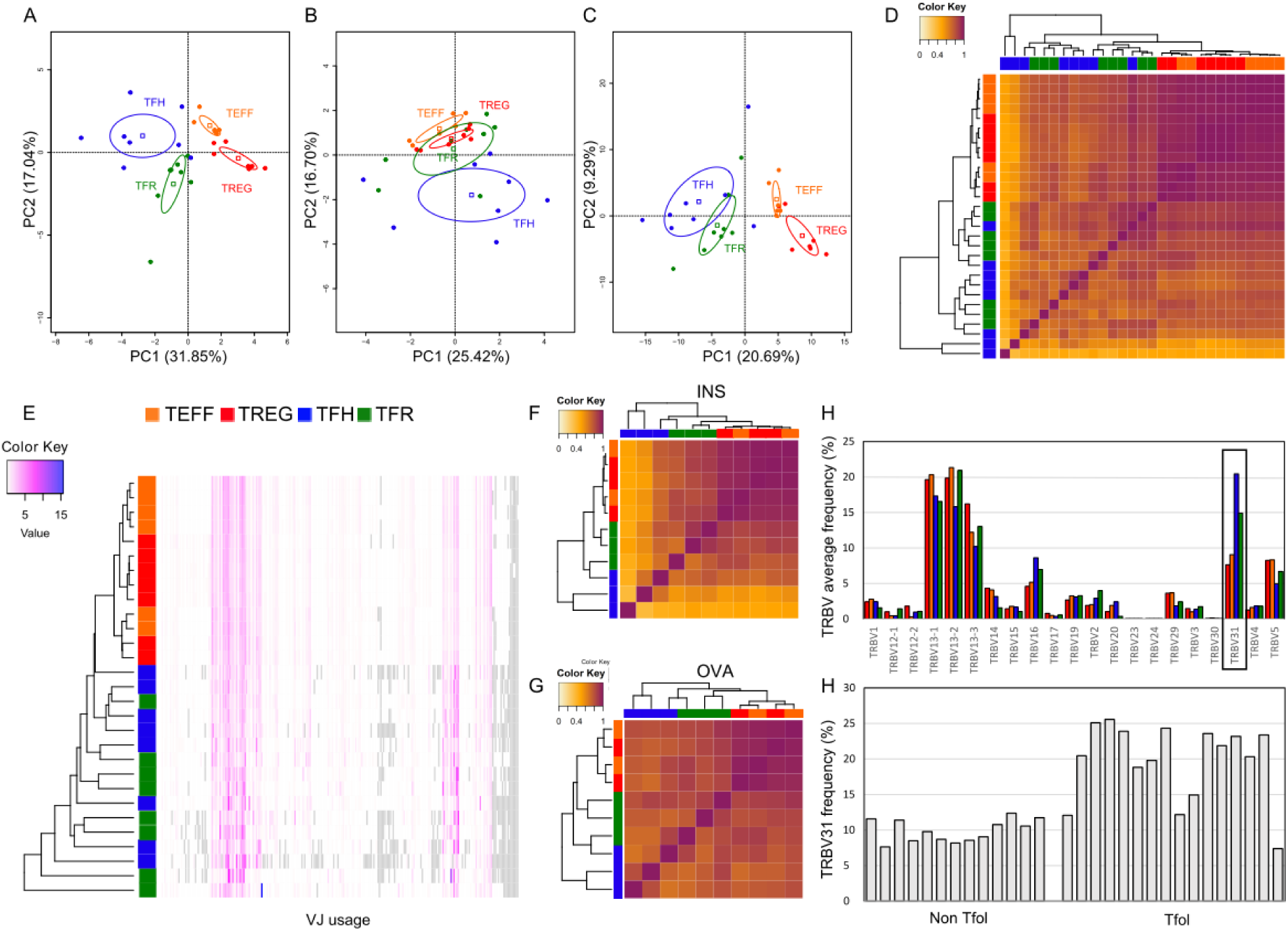
TRBV gene usage separates Tfol from non-Tfol cells. (**A-C**) PCA projection of the four subset samples according to the first two components (x-axis:PC1; y-axis:PC2) is plotted for the TRBV usage (**A**), TRBJ usage (**B**) or TRBVBJ usage (**C**) of the samples. (**D**) Hierarchical clustering heatmap of Morisita-Horn similarity index values for all pairs of samples according to the indicated colour scale. (**E**) Hierarchical clustering of TRBVBJ frequencies across samples. **(F-G)** Same analyses as D but for individuals immunized with OVA (**F**) or INS (**G**). (**H**) Bar plot showing TRBV gene usage for four representative samples of the four T cell subsets. (**I**) Bar plot showing the TRBV31 usage among all non-Tfol vs. Tfol samples.

We next computed the Morisita—Horn (MH) index between all samples based on their TRBVBJ usage (**Fig. 2D**), which calculates a similarity score ranging from 0 (dissimilar) to 1 (similar) between each pair of samples, and performed a hierarchical clustering (Euclidean distance and “complete” method) of all the samples using this metric. In line with the PCA analysis, Tfh- and Tfr-cell samples (Tfol) were clustered together, apart from the Teff and Treg (non-Tfol) cells. However, Tfol samples are intermingled within their cluster. Although their TRBV and TRBVBJ usage appeared to distinguish regulatory vs. helper cells on PCA for Tfol and non-Tfol cells, the sharing of TRBVBJ gene usage between Treg and Teff cells is higher than that between Tfh and Tfr cells (**Fig. 2D**, **Supplementary Fig. S2**). Hierarchical clustering of TRBVBJ frequencies (**Fig. 2E**) also showed Teff and Treg cell co-clustering. In contrast, Tfol cells do not cluster together but are just separated from non-Tfol cells. This suggests heterogeneity among Tfol-cell samples.

Since samples originated from mice undergoing different immunization protocols, we independently computed the MH similarity matrix between samples from mice immunized with either INS (**Fig. 2F**) or OVA (**Fig. 2G**). Similar observations were made.

In order to understand whether the difference of TRBV and TRBVBJ usage between samples on PCA was due to major changes in TRBV usage at the individual level, we plotted the frequencies of TRBV genes of the four subsets and observed no major differences except for TRBV31 gene expression, which was overexpressed in Tfol cells (**Fig. 2H**). We confirmed these observations in all our samples (**Fig. 2I**). This further demonstrates a peculiar TRBV repertoire in Tfol vs. non-Tfol samples.

### The clonotype distribution is different among the four cell populations

We further analyzed the data by exploring the diversity at the clonotype level. The projection of clonotype frequencies by PCA was performed on clonotypes shared by at least five samples to reduce noise due to private clonotypes. Tfol cells are well separated from non-Tfol cells on PC1 (34%). Strikingly, in contrast with our observations on the TRBVBJ usage, we observed that Tfh and Tfr cells are remarkably close to each other, while Teff and Treg cells are rather well separated (**Fig. 3A**). When focusing on Tfh and Tfr cells only, their differences were revealed. The two subsets are separated on the first two components with a PC1 of 26%, suggesting that the two subsets are close in Figure 3A because they have similar overall characteristics compared to non-Tfol cells, but are in fact different at a closer clonotype level (**Fig. 3B**). As expected, PCA also well separated Treg and Teff cells (**Fig. 3C**).

**Fig. 3:**
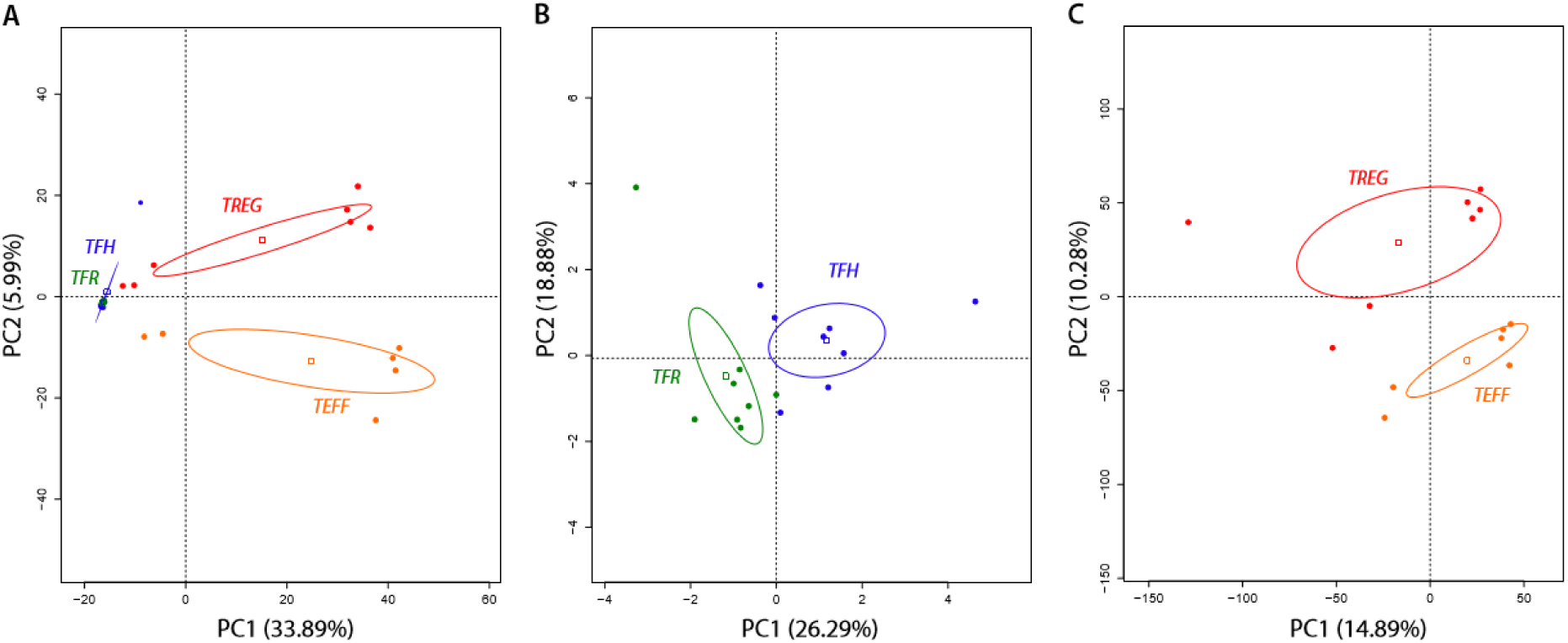
Clonotype composition distinguishes Tfr from Tfh cells. (**A-C**) PCA is plotted according to the first two components (x-axis:PC1; y-axis:PC2) using the frequencies of the predominant clonotypes shared by at least five samples across the Tfr, Tfh, Treg and Teff cells (**A**), Tfr and Tfh cells (**B**) and Treg and Teff cells (**C**).

### Tfr and Tfh cells have distinct repertoires

We first compared the clonotype composition of Tfr and Tfh cell repertoires irrespective of the immunization. We ordered all clonotypes of a given Tfr-cell sample by decreasing frequency and selected the 250 predominant clonotypes. For each of the 8 Tfr samples, we evaluated the frequencies of these 250 clonotypes in each of the seven other Tfr cell (or Tfh cell) samples and calculated a mean frequency of these 7 values (**Fig. 4A**). We plotted these means for each of the eight available samples (**Fig. 4B**). We used the same methodology with Tfh cell predominant clonotypes (**Fig. 4C**). Results showed that, irrespectively of immunization, clonotypes found predominantly in Tfr cells are also found in higher proportions in other Tfr-cell samples than in Tfh-cell samples (**Fig. 4B**). Conversely, clonotypes predominantly found in Tfh cells are mostly shared with other Tfh-cell samples rather than with Tfr-cell samples (**Fig. 4C**).

**Fig. 4:**
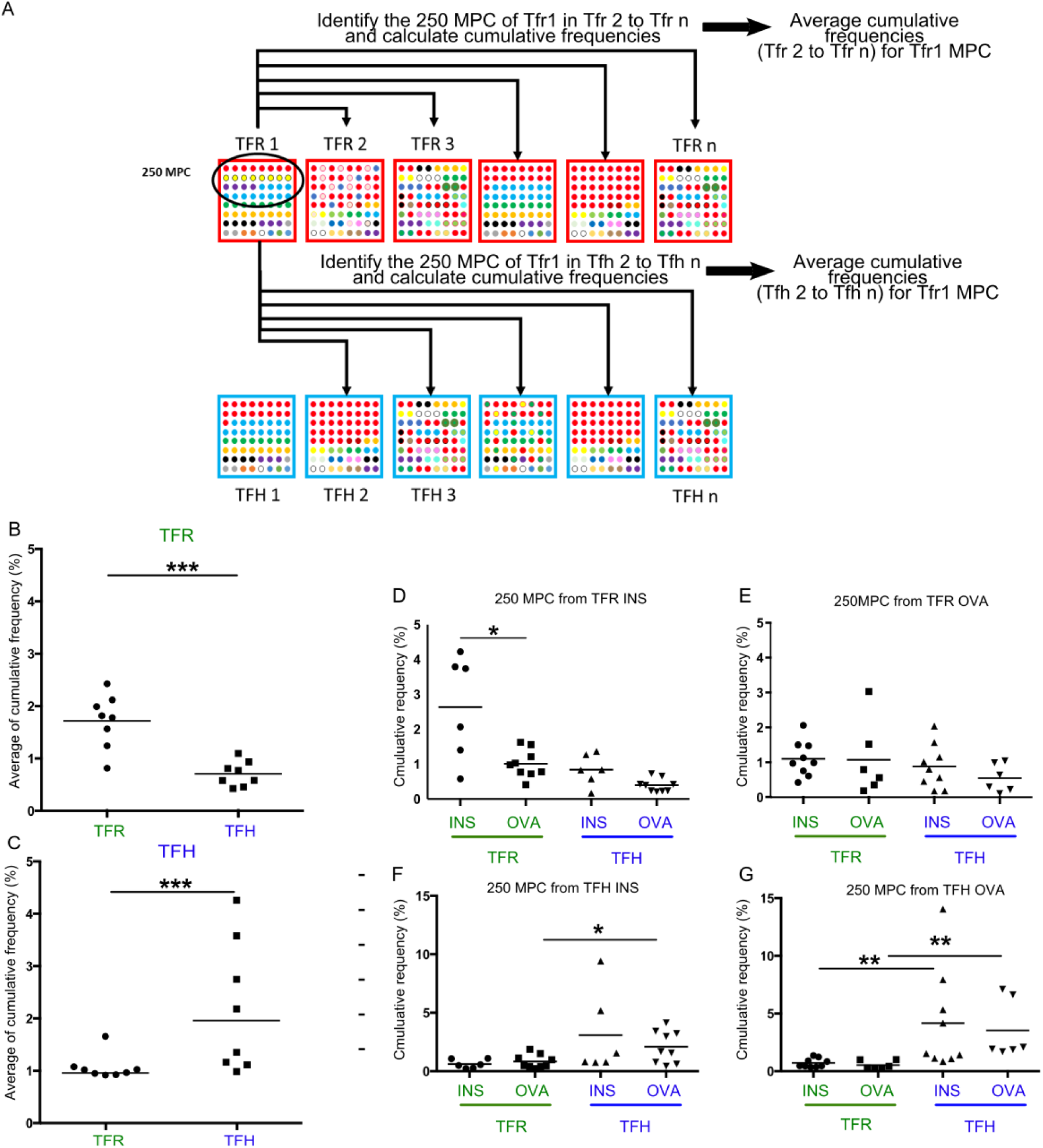
Tfr- and Tfh-cell predominant clonotypes are unique to each subset. (**A**) Representative illustration of the average cumulative frequencies calculated for the 250 most predominant clonotypes (MPC) of Tfr (**B**) and Tfh (**C**), (**B-C**) Plots showing the average of the predominant Tfr-cell clonotypes among Tfr- and Tfh-cells samples of the other seven individuals (**B**), and of predominant Tfh-cell clonotypes among Tfr- and Tfh-cell samples of the other seven individuals (**C**). (**D-G**) Plots showing the representation of the predominant clonotypes from Tfr cells (**D-E**) and Tfh cells (**F-G**) from INS (**D, F**) and OVA (**E, G**) immunized mice among Tfr- and Tfh-cells samples of the other seven individuals depending on the immunizing antigen.

We then performed a similar analysis for comparing the response to specific immunization. The 250 most predominant clonotypes of each of the Tfr cell samples from mice immunized with INS were analysed (i) in the two other Tfr samples of INS immunized mice, generating 6 values that were plotted individually; and (ii) in the 3 other samples from Tfh from INS immunized mice, or the Tfr and Tfh of OVA immunized mice, generating 9 values per comparison, plotted individually (**Fig. 4D**). A similar analysis was performed for Tfr samples from OVA immunized mice (**Fig. 4E**). Results showed that frequent clonotypes from Tfr cells of mice immunized with INS are found in higher proportions in other Tfr-cell samples from mice immunized with INS than in the other Tfr-cell or Tfh-cell samples (**Fig. 4D**). This phenomenon was not observed for Tfr cells of mice immunized with OVA (**Fig. 4E**) suggesting that within the global repertoire a Tfr-self-antigen specific response can be detected. Conversely, Tfh cell major clonotypes were similarly found in Tfh cells regardless of the immunizing antigen (**Fig. 4F-G**), suggesting that an antigen specific Tfh response cannot be detected within the global repertoire.

We further analysed the specific response to immunization within a more restricted repertoire of public clonotypes, i.e. those shared by three similar samples. Public clonotypes from Tfh samples of OVA-immunized mice represent an average of 5% of the repertoire of these cells, but less than 1% of the repertoires of the other categories of cells (Tfr cells from all conditions or Tfh cells from mice immunized with INS or nothing) (**Fig. 5A**). In contrast, public clonotypes from Tfh cells of mice immunized with INS were equally represented among all samples, amounting to 2% of their repertoire (Tfh and Tfr cells, irrespective of the immunization) (**Fig. 5B**). Conversely, for Tfr cells there was three-fold higher clonotype sharing between Tfr cells from mice immunized with INS than for mice immunized with OVA (**Fig. 5C-D**).

**Fig. 5:**
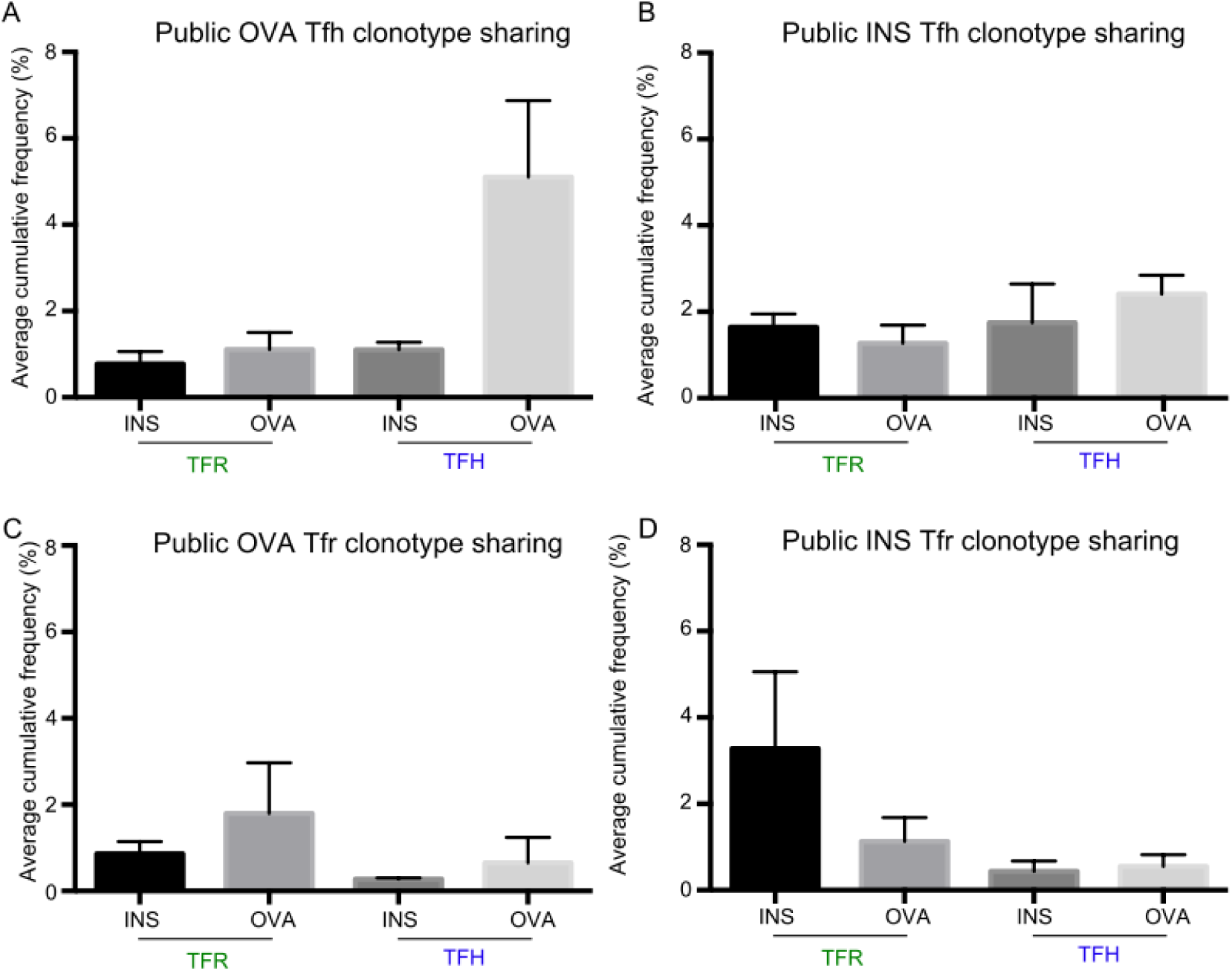
Tfr and Tfh are respectively self and non-self-antigen specifics. (**A-D**) Histograms showing the averages of cumulative frequencies of predominant clonotypes shared by the three Tfh-cell subsets across all subsets from mice immunized with OVA (**A**) or INS (**B**), or shared by the three Tfr subsets across all subsets from mice immunized with OVA (**C**) or INS (**D**).

Finally, we then attempted to reveal similarities between repertoires at a less stringent level using “grouping of lymphocyte interactions by paratope hotspots” (GLIPH) that clusters TCRs with a high probability of recognizing similar antigens owing to either conserved motifs of 2 to 4 amino-acids and/or global similarity of complementarity-determining region 3 (CDR3) sequences(31). The majority of motifs shared by the 3 Tfr samples of mice immunized by INS were found into those shared by Tfr samples from mice immunized by OVA or from nonimmunized mice (**Fig. 6A**). The same observation was made for Tfh cells (**Fig. 6B**).

**Fig. 6:**
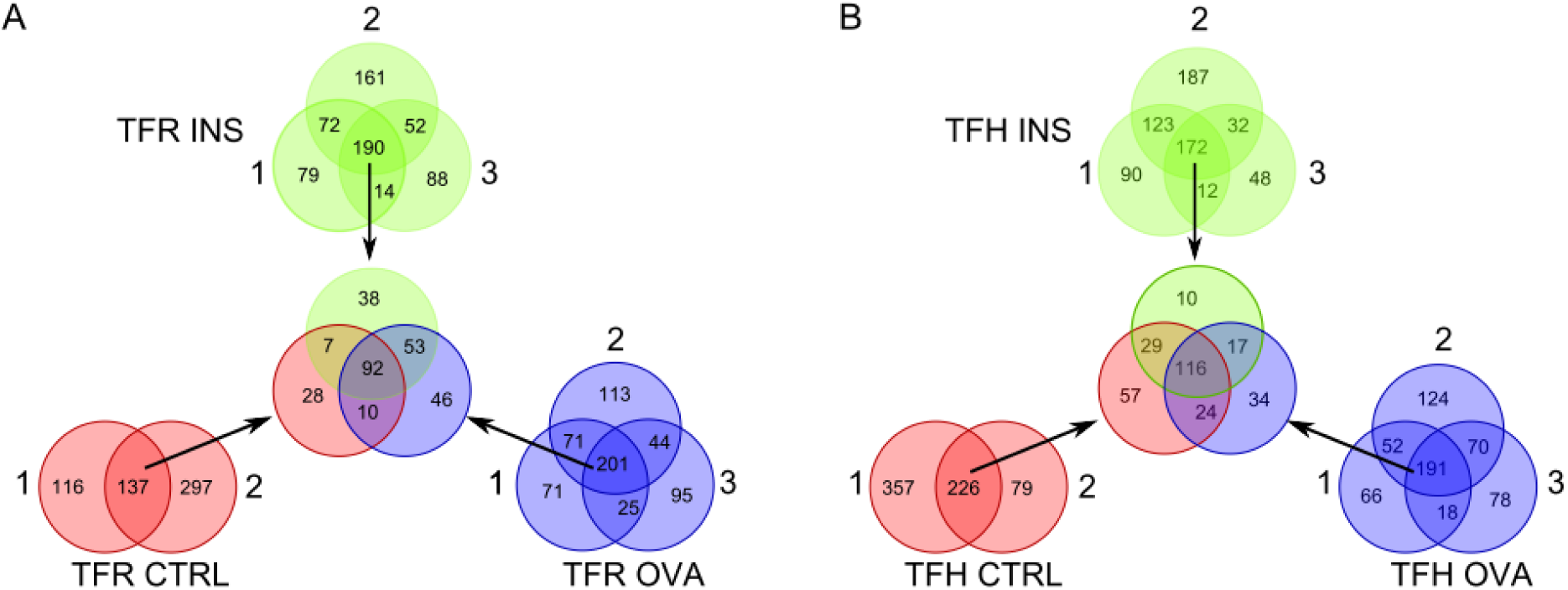
Tfr- and Tfh-cell predominant motifs are independent of the immunizing antigen. Venn diagrams showing shared motifs between Tfr-cell samples (A) and Tfh-cell samples (B). First, public motifs from Tfr (A) of non-immunized mice (red) or mice immunized with INS (green) or OVA (blue) were identified and then compared between each othet. Same analysis was performed for Tfh motifs (B).

### Tfr cells share more of their repertoire with Treg than with Teff cells

We represented the sharing of major clonotypes using a Venn diagram. The sharing of the 1% predominant clonotypes of Tfh, Tfr, Treg and Teff cells revealed that 13% of Tfr cell clonotypes are shared with Treg cells, while only 6% are shared with Teff cells. Conversely, 1,7% of Treg cell clonotypes are shared with Tfr cells, while only 0.8% are shared with Tfh cells (**Fig. 7A**). These observations are confirmed by the analysis of clonotypes sharing at the individual level. Those of the 250 predominant clonotypes of Treg cells present in Tfr or Tfh cells represent an approximately 5 times higher percentage of the repertoire of Tfr than Tfh cells, with numerous expanded clonotypes (**Fig. 7B-C**). Similarly, those of the 250 predominant clonotypes of Tfr cells present in Treg or Teff cells represent a higher percentage of the repertoire of Treg than Teff cells (**Fig. 7D-E**).

**Fig. 7:**
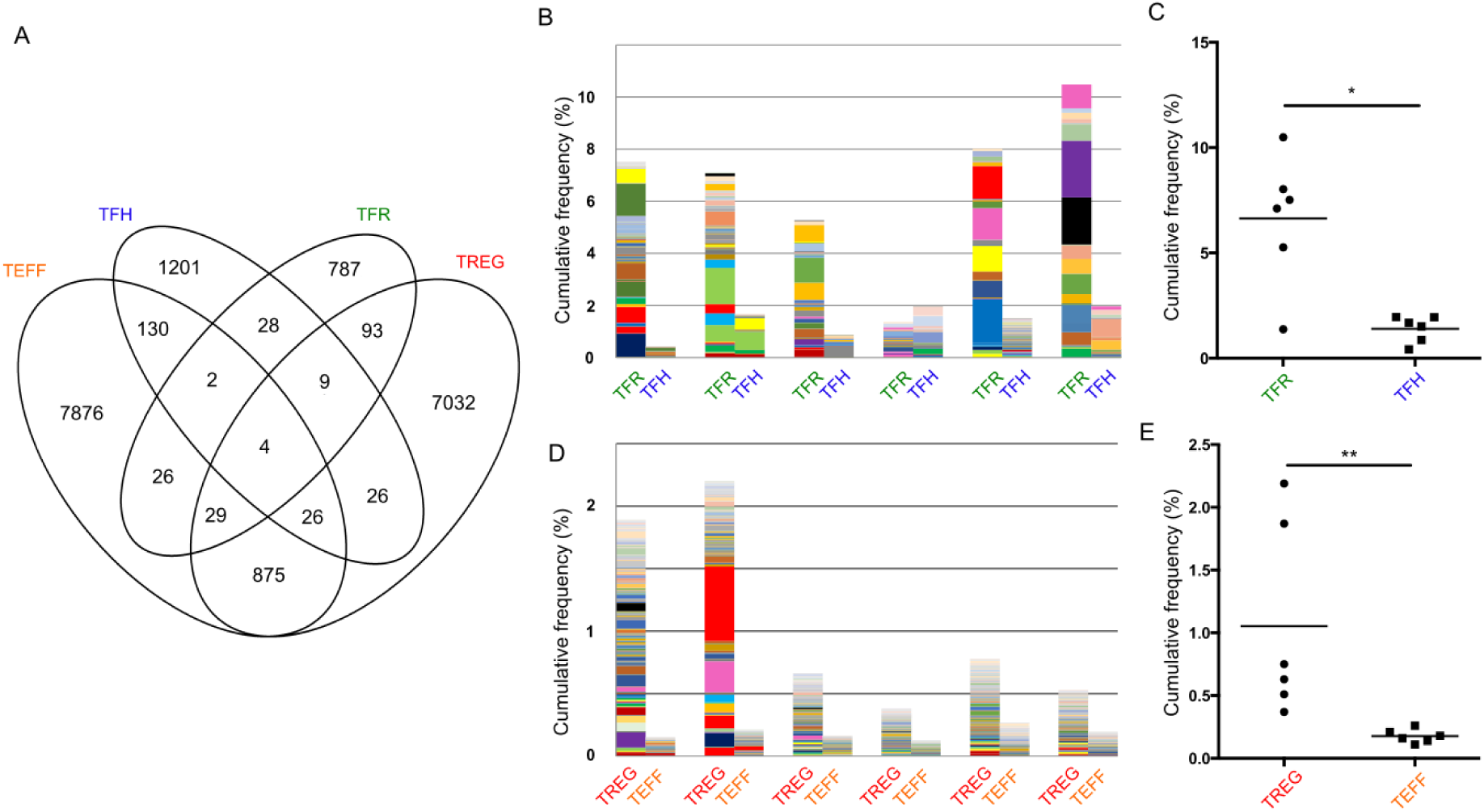
Tfr-cell repertoire supports a Treg-cell origin. (**A**) Venn diagram between the 1% predominant clonotypes of Tfh, Tfr, Treg and Teff cells. (**B**) Histograms showing the cumulative frequency of the predominant Treg clonotypes shared with Tfr and Tfh cells. (**C**) Histograms showing the cumulative frequency of the predominant Tfr clonotypes shared with Treg and Teff cells. (D-E) Statistical analysis of the histograms shown in B (**D**) and C (**E**) (Mann-Whitney U, *: p<0.05; **: p<0.01, ***: p<0.001).

## DISCUSSION

### Tfh and Tfr cells have a higher TCR diversity than expected

Tfol cell TCR repertoires are less diverse than those of non-Tfol cells (**Fig. 1**), but still surprisingly diverse. Indeed, these cells that expand in response to immunization are stringently identified (20) by markers that assign them to GCs, a specialized site in which antigen-specific antibodies are formed (2). In the GCs, antigen-specific B cells act as antigen-presenting cells for Tfh cells, implying that the B cells and the Tfh cells are specific for the same antigen. It could thus have been conjectured that Tfh cells that are responding to an immunization would have a repertoire limited to a few clonotypes, with large expansions. Instead, we found thousands of sequences in every Tfh- and Tfr-cell sample (**Fig. 1**), a point that was missed by analyzing Tfh cells purified using tetramers (28) or from mice bearing a fixed TCRβ chain (29). This observation indicates major bystander Tfol activation during the response to immunization. The number of Tfh cells increase approximately by 10 times after immunization with OVA. Assuming 40 main epitopes for OVA (32) and 1/40 000 the frequency (33) of T cells responding to each of these, this Tfh cell expansion should have resulted from an approximate 9000 fold expansion of these antigen specific cells, leading to mostly detect these much expanded clonotypes. This is clearly not observed, as the predominant 1% of clonotypes for Tfh cells represents approximately only 50% of the total repertoire.

Thus, our results establish that the overall expansion of Tfh cells after immunization results in part from the expansion of cells that are not responding to main OVA epitopes (nor to proteins from the adjuvant as we used alum). This suggest that the initial trigger of the immune response provided by dendritic cells could not only activate high-affinity antigen-specific Tfh cells (25), but also bystander T cells with specificity for other antigens presented by these dendritic cells and/or activated under the effects of the cytokine environment. In this line, this bystander activation could be related to the major role of IL-1 in Tfh-cell activation that we recently reported (20). Alternatively, or complementarily, the very high cross-reactivity of single TCR that have been shown to recognize thousands of different peptides (33) could explain the bystander activation of cells bearing promiscuous TCRs with low affinity for the immunizing antigen, thus less strongly stimulated and not driven to highly proliferate. Looking at motif sharing further supports this as we found that a majority of motifs are shared between Tfr (or Tfh) cells regardless of the immunization.

Similarly, our results show that Tfr-cell repertoire diversity is high, regardless of the nature of the immunizing antigen. This bystander effect for Tfr activation is less unexpected as even in the context of a specific immunization, many other self-antigens are expressed by APCs.

### Tfh-cells predominantly respond to foreign antigens while Tfr-cell respond to self-antigens

At the clonotype level, we confirmed previous observations (29) that Tfh and Tfr cells have two distinct repertoires. The predominant clonotypes of Tfr cells in one individual are systematically found at higher frequencies in Tfr cells of other individuals than in Tfh cells, and inversely for Tfh-cell predominant clonotypes.

Sharing of predominant clonotypes (either public or not) indicate that the Tfh cell response is mostly towards foreign antigens while the Tfr cells response is mostly towards self-antigens. Mirrored observations were made for Tfr cells that appear to respond to self-rather than foreign-antigens. This is in agreement with our recent observation that Tfr cells increase significantly more in INS-compared with OVA-immunized animals (20). Therefore, the Tfr repertoire in OVA-immunized animals could reflect a bystander recruitment of Tfr cells, independently of the immunizing antigen, while the Tfr repertoire in INS-immunized mice could also comprise a response to the immunizing antigen. This is also in line with a recent study showing that Tfr cells have an effect only on self-reactive antibody responses (34)

### The Tfr-cell repertoire is close to the Treg-cell repertoire

Using stringently purified Tfr cells, devoid of contaminating Treg cells, we show that clonotype sharing likens the Tfr cell repertoire to the Treg-cell repertoire (**Fig. 7**). This provides an indication of the origin of Tfr cells. Indeed, it should be more difficult to mimic a repertoire than a phenotype. The follicular phenotype has been shown to be mainly dependent on the expression of Bcl6, and is thus triggered by the expression of this unique molecule (7, 35, 36). In contrast, major sharing in a repertoire is dependent on many complex processes, from cell selection in the thymus to the dynamics of post-thymic cells. Thus, the fact that two subsets (here Tfr and Treg cells) share more of their repertoire than with the other two subsets (Tfr and Tfh cells) is indicative of a higher probability of a common origin.

Altogether, our observations made for the first time with the full repertoire of *bona fide* CD25^−^ Tfr cells that are not contaminated by Treg cells (20),(21,22) highlight a repertoire related to that of Treg cells and as such that is in large part self-specific (37, 38) and a bystander activation of these cells.

This bystander effect could be evidenced because follicular cells (i) are terminally differentiated cells found in a highly specialized structure that should contain only cells responding to the immunization, and (ii) can be unambiguously identified and purified. While we are often biased to detect and focus on antigen specific responses, we believe that such a bystander activation could be a more general phenomenon in the T cell response to antigens. It is now becoming clearer that in addition to specificity, there is an extraordinary fuzziness and plasticity in the immune system, which needs to be taken in consideration. Actually, autoantibodies have been found in healthy humans and mice in the absence of an immunization with their target antigens (39). It thus remains to study whether the bystander activation of Tfol has relevance for the development of immunopathologies.

## Material and Methods

### Mice

8- to 14-week-old male and female NOD Foxp3-gfp mice, which express the green-fluorescent protein (GFP) under the control of the Foxp3 gene promoter, were provided by V. Kuchroo, Brigham and Women’s Hospital, Boston, MA. All animals were maintained at the University Pierre and Marie Curie Centre d’Expérimentation Fonctionnelle animal facility under specific pathogen-free conditions in agreement with current European legislation on animal care, housing and scientific experimentation (agreement number A751315). All procedures were approved by the local animal ethics committee.

### Immunization

Mice were immunized once (D0) and sacrificed at D10. Intraperitoneal injection was performed with 100 μg of OVA (Ova A5503; Sigma-Aldrich) mixed with 500 μg of aluminium hydroxide (alum) gel (AlH303; Sigma) or with 4.5 IU of human insulin (Umuline Rapide; Lilly) mixed with 500 μg of alum.

### Cell sorting

Splenocytes from immunized mice were stained with Ter-119-biotin and B220-biotin antibodies for 20 min at 4°C and labelled with anti-biotin magnetic beads (Miltenyi Biotec) for 15 min at 4°C. B cells and erythrocytes were depleted on an AutoMACS separator (Miltenyi Biotec) following the manufacturer’s procedure. Enriched T cells were stained for 20 min at 4°C with the following monoclonal antibodies at predetermined optimal dilutions: CD4-V500 (BD Biosciences), CD8a-AF700 (BD Biosciences), streptavidin-APC (eBioscience) or -APC-Cy7 (BD Biosciences), PD-1-PE (eBioscience), CXCR5-Biotin (BD Biosciences). CXCR5 staining was performed using biotinylated anti-CXCR5 for 30 min at 20°C followed by APC- or APC-Cy7-labelled streptavidin at 4°C. The following subsets were sorted on a BD FACSAria II (BD Biosciences) with a purity > 98%: CD4^+^CD8^−^CXCR5^hi^PD-1^hi^Foxp3^−^ T follicular helper T cells (Tfh); CD4^+^CD8^−^CXCR5^hi^PD-1^hi^Foxp3^+^ follicular regulatory T cells (Tfr); CD4^+^CD8^−^ CXCR5^int/lo^PD-1^int/lo^Foxp3+ regulatory T cells (Treg) and CD4^+^CD8^−^ CXCR5^int/lo^PD-1^int/lo^Foxp3^−^ effector T cells (Teff). Sorted cells were stored in lysis buffer (Ambion) at ‒80°C until processing.

### TCR deep sequencing

RNA from sorted cells was extracted using the RNAqueous kit (Ambion) and sent to iRepertoire^®^ (Huntsville) for cDNA synthesis and TCR amplification following their protocol (40). Briefly, each TCR is reverse-transcribed using a set of 24 forward primers, each targeting one TRBV mouse gene, and a reverse primer located in the TRBC gene to ensure a complete coverage of the clonotype sequence. PCR1 primers include barcodes to allow sample identification after the sequencing. PCR1 products are then purified on magnetic beads and a second round of PCR is performed. Amplified libraries are excised from agarose gel and purified. Paired-end sequencing is then carried out on a Miseq Illumina sequencer using a 2x250-bp read length protocol.

### TCR deep sequencing data processing

Each FASTQ raw data file obtained from iRepertoire was processed for TRB sequence annotation using the clonotypeR toolkit and associated R packages (41). Each dataset can be summarized as a list of clonotypes (defined as a unique combination of TRBV-CDR3aa-TRBJ sequence) and their associated counts in the dataset. These values are computed to quantify the differences between repertoires at several complementary levels. Clonotypes observed only once in a dataset were discarded.

### Data analysis

Statistical comparisons and multivariate analyses (PCA, hierarchical clustering, Venn diagrams) were performed using R software version 3.1.3 (www.r-project.org). The Morisita-Horn index (42) assesses the similarity between sample sets. It ranges from 0 (no common species between the two samples) to 1 (all species are equally present in the two samples). Unlike the Morisita index, the Morisita-Horn variant takes into account the relative abundance of species in the sample. P50 was calculated as the percentage of unique predominant clonotypes necessary to reach 50% of the total number of sequences in a given sample. Conserved motifs and global similarity of the CDR3 was assessed by “grouping of lymphocyte interactions by paratope hotspots” (GLIPH) (31) on the 10% most predominant clonotypes per sample. Motifs were first identified for each cell subset and per experimental condition. The motif sharing was assessed per experimental condition. Then the motifs shared by all the samples from a given experimental group were compared with those shared by all the samples from each of the other expiremental condition per cell subset.

Statistical analysis were performed using the non-parametric Mann-Whitney U test on GraphPad Prism v5 (p-values are indicated in the figures such as ns: p > 0.05, *: p < 0.05, **: p < 0.01, ***: p <0.001).

## Acknowledgments

We are grateful to B. Gouritin and F. Brimaud for help in cell sorting and to G. Churlaud for help in experiments. We thank V Kuchroo for the mice provided.

## Author contributions

PGR and BB performed the mouse experiments. PGR analyzed all the mouse data. KES performed the GLIPH study. PGR, EMF and DK conceived the experiments. WC conceived the workflow of analysis. PGR, EMF, AS and DK wrote the manuscript with input from all authors. DK conceived, supervised and obtained funding for the entire study.

## Data and materials availability

All the data can be made available upon request.

